# The Creative Mind: Blending Oxytocinergic, Dopaminergic and Personality

**DOI:** 10.1101/700807

**Authors:** Anne Chong, Benjamin Becker, Dario Cruz Angeles, Maria Gutierrez Matos, Xiong Yue, Poh San Lai, Mike Cheung, Zhen Lei, Fabio Malavasi, Soo Hong Chew, Richard P. Ebstein

**Author notes:** To whom correspondence should be addressed author Richard P. Ebstein, C2BEF (China Center for Behavior Economics and Finance), Southwestern University of Finance and Economics (SWUFE), Chengdu, China.

## Abstract

In a fast-changing world, creative thinking (CT) is an extraordinary currency. Oxytocin (OT) is associated with CT and release of OT depends on ADP ribosyl-cyclases (*CD38* and *CD157*). Moreover, CT as well as OT’s mechanism of action are mediated via central dopaminergic pathways. Consequently, we evaluate the roles of *CD38*, *CD157*, dopamine receptor D2 (*DRD2*) and catechol-O-methyltransferase (*COMT*) peripheral gene expression in CT. Two principal domains of CT, divergent thinking and insight solving problems, were assessed using validated behavioral assessments. To facilitate discriminant validity, two established correlates of CT, trait Openness and fluid intelligence as well as age and sex were included in the regression model. In women, significant main effects (p<0.01) were positively associated with the expression *CD38*, *CD15*, and their interaction *CD38* x *CD157* controlling for Openness, fluid intelligence and age. Subsequent analysis on the subscale-level revealed significant main effects for *CD157* and *CD38* x *CD157* in men specifically for divergent thinking. In women, significant (p<0.01) results are also observed for dopaminergic expression (*DRD2*, *COMT*, *DRD2* x *COMT*). The full model (oxytocinergic and dopaminergic gene expressions, Openness, and fluid intelligence) explains a sizable 39% of the variance in females. Significant main effects are observed for *CD38*, *CD157*, *DRD2* and *COMT* as well as their interactions (*CD38*x*CD157* and *DRD2*x*COMT*). In conclusion, we show that oxytocinergic and dopaminergic gene expression contribute significantly to the complex CT phenotype suggesting the notion that the perspective gained from examining the peripheral transcriptome meaningfully adds to understanding the landscape of creative thinking.

**Significance Statement:** Creative thinking (CT) is a powerful driving force galvanizing progress and civilization. Towards better understanding the neurobiology of CT, we implement a gene expression strategy that is considered to capture not only genomic elements but also environmental signatures. We employ laboratory measures of CT (alternative uses test and insight problem solving), controlling for fluid intelligence and the personality trait of Openness. We focus on oxytocinergic and dopaminergic genes that contribute to the molecular architecture of CT. Oxytocinergic and dopaminergic gene expression significantly explains a robust 39% of the variance in CT. Notably, this study demonstrates the potential of the peripheral transcriptome towards tracing gene pathways underlying some complex cognitive behaviors such as creative thinking.

## Introduction

Creativity underpins the advancement of civilization and has driven the progress of H. *sapiens* from the middle Pleistocene to the current post-industrial Information Technology age. In ancient times, creativity was thought to be a gift of the divine, viz. “and I have put wisdom in the hearts of all the gifted artisans, that they may make all that I have commanded you: the tabernacle of meeting…” Exodus 31-32. For the past 2000 years there have been shifts in understanding creativity, from Aristotle, who first associated creativity with madness, to Galton who replaced the divine with an empirical approach by studying genealogies characterized by creative genius (1). The modern era of creativity research can be pegged to the address by Guilford, then President of the APA in 1950 (2). Among the most widely-applied definitions of creativity, the requirement for ideas that are both original and useful or appropriate stands out (3). Creativity is a complex behaviour that is characterized by marked individual differences in cognitive flexibility and ability (exemplified by divergent thinking and problem solving) (4) and open-mindedness, assessed in the Big Five personality trait of Openness to experience (5), a core trait underlying creativity (6). Creativity, not surprisingly, is also underpinned by fluid intelligence (7).

There is some evidence that creativity is in part due to heredity. Early small twin studies reveal a modest heritability of around 20%, with the majority of the variance assigned to shared and nonshared environmental effects (8). A more recent analysis shows that the genetic variance in tested figural creativity is explained in part by the genetic variance in intelligence and the two personality trait of Openness (3). It appears that personality traits and intelligence are partial mediators of the genetic effects on individual differences in creativity. Somewhat higher levels of heredity are observed using other measures of creativity. Moderate to high heritability was observed in a large twin study that focused on subjects reporting a creative profession as the phenotype (9). Like other complex traits, additive genetic effects and unique environmental factors play the major roles. Multiple measurements of creative achievement suggest a heritability of 43-67% and the remaining variance is due to nonshared environmental influences (10).

The personality trait of Openness to experience is one of the most robust contributors to CT with correlations ranging between 0.2 to 0.5, see (6) among many. Individuals high in Openness to experience are broadminded, creative, curious, and cultured. There are several avenues by which enhanced Openness can boost creativity. Openness encompasses traits such as vivid imagination and intellectual curiosity, the ability to entertain unfamiliar and unconventional thoughts, and use combinatory processing with previous knowledge. Therefore, high Openness can seed and accelerate overall creativity. A second important contributor to creativity is intelligence. In particular, enhanced fluid intelligence allows the use of more complex information garnered from ones’ surroundings, speedier information processing, and a more critical evaluation of the suitability of new ideas (11). Many studies point to an important role of intelligence in creativity (12). A recent study examined predictors for real-life creative outcomes (13) and found Openness to experiences and creative potential assessed by divergent thinking (DT) using the alternative uses test (AUT), predict everyday creative activities. Creative activities then predicted real-life creative achievement. The same study found intelligence predicts creative achievement, but not creative activities. By all accounts, creativity is a multifaceted construct in which heredity and environment play a joint role in contributing to creative thinking and real-life achievements.

Several studies have tested specific polymorphic genes for a role in creativity (14, 15). These candidate gene studies, albeit small, especially highlight two neurotransmitter systems in contributing to creativity: oxytocinergic and dopaminergic pathways. Of especial interest with respect to oxytocin (OT) are findings of De Dreu and colleagues (15, 16) suggesting that, in experiments using intranasal oxytocin (OT) administration and testing SNP associations within the oxytocin receptor gene (*OXTR*) region, make a case for a role of OT in contributing to creativity. They review the literature and find mixed evidence for a role of *OXTR* SNPs in creative thinking (16). Interestingly, they also examined *CD38*, and in three out of the four investigations, major frequency allele carriers of *CD38* SNPs scored lower on neuroticism and higher on imagination, personality traits related to creativity. CD38, is a type II transmembrane glycoprotein with ADP-ribosyl cyclase activity located both peripherally and in the brain (17) that governs central release of OT (18). CD38 has numerous purposes acting as receptor, ectoenzyme (19) and second messenger playing the role of a ubiquitous calcium-signalling molecule (20). CD38 is also a well-studied immune cell marker whose expression increases in some diseases such as chronic lymphocytic leukaemia (CLL) (21). Dopamine (DA) has been long considered to contribute to CT and two widely-studied polymorphic DA genes have been specifically linked to creativity, *DRD2* (14) and *COMT* (22). Mesocortical DA projections to the forebrain are known to be involved in cognition (23) and hence are also likely important in CT (24). Notably, accumulating evidence suggests that these pathways are modulated by both, DA as well as OT (25, 26) and that the two systems interact with respect to the multi-facetted behavioural roles of OT (e.g. (27)

Against this background the current investigation examines the role of oxytocinergic and dopaminergic pathways in contributing to creativity by using laboratory-based assessments of the two principal domains of creativity, divergent thinking (DT) measured by the alternative uses test (AUT) (28) and insight solving problem (IPS) (29). IPS often gives rise to the so-called “Aha!” or Eureka moment. AUT is one of the most widely used assessments of domain-general DT (30). Guilford (31) defined four components of DT that are evaluated by the AUT (i.e. fluency, flexibility, originality and elaboration), with originality – defined as novelty or infrequency of ideas – being considered as “vital for creativity” (Runco et al (32)) and matching the originality facet of the CT conceptualization as “original (and useful or appropriate)” (33). Along these lines the use of originality as a measure of novelty in the AUT is well-supported (34).

Insight problems require solutions that are apparently spontaneous and often attributed to unconscious, associative processes (35). Successful IPS problem-solving requires restructuring the problem and “thinking outside the box” (36). IPS has been correlated to the convergent thinking facet of creativity (37) where distally related concepts are brought together towards a novel solution. CT is a complex cognitive behavior and as such inherently associated with broader personality dimensions, specifically Openness (5) and underlying general cognitive abilities such as fluid intelligence (38) which serves as basic building blocks of several higher order cognitive domains. To provide an important benchmark and increase the discriminant validity (39) of the laboratory measures of CT, as well as to avoid critically confounding effects of intelligence and Openness (see also (40)), the personality trait of Openness (5) measured using the NEO-PI-3 and fluid intelligence (38) indexed by the short version of Raven’s Standard Progressive Matrices (41) were additionally assessed in the present study.

Towards examining the role of neural gene pathways in contributing to CT (AUT and IPS), we examine gene expression for oxytocinergic (*CD38*, *CD157*) and dopaminergic (*DRD2, COMT)* gene expression in saliva samples and including Openness, fluid intelligence, and sex in our model. Many effects of oxytocin are sex dependent, e.g. (42–50) perhaps suggesting that the OT system might have evolved differential roles in men and women, for instance oxytocin’s function in mammals may be especially crucial to female reproductive behavior, viz. helping regulate parturition and lactation. It seems reasonable, therefore, that sex would strongly bear on *CD38* and *CD157* gene expression.

*CD38* and its homologue *CD157* (BST-1), contiguous gene duplicates on human chromosome 4 (4p15), represent a gene family that regulates cellular interactions (19). Importantly, an oxytocin analogue has long lasting effects on anxiety behavior in a *CD157* knockout mouse (51) and communication impairment during the suckling period is restored by oxytocin in the *CD157* knockout (52). *CD157* and *CD38* knockouts show decreased plasma oxytocin levels (53) suggesting a role for both, *CD157* as well as *CD38* in oxytocin (OT) release and thus as putative regulators of complex behaviors (53). Consequently, we chose to examine both homologues in the current study. A number of studies show that saliva is a reasonable target tissue for gene expression studies (54) including determination of alterations in behavioral disorders (55). The use of peripheral biomarkers, and gene expression in particular, to better understand the combined role of genes, their pathways and environment, is gaining traction (56) in clinical psychiatry (57), neurology (58), drug discovery (59), cancer treatment (60) and psychology (61). Saliva has been characterized as the “mirror of the body” and the “perfect medium to be explored for health and disease surveillance” (62). Importantly, technological strides enable stabilization of salivary RNA for downstream genomic applications (63).

Crucially, gene expression captures both genetic and environmental information and inherently appears to have greater predictive power than the current genome-wide association studies (GWAS) based exclusively on analysis of single SNPS, which account for only a tiny percent of the variance, e.g. 0.02% (64, 65). Moreover, gene expression studies can be robustly coupled with other predicting and controlling variables (66), as shown in the current analysis of CT, and notably these models regularly explain a considerable percentage of individual differences for complex phenotypes.

## Results

### Descriptive Statistics

Table 1 shows the descriptive statistics of the study participants (n = 145) and the various measures including *CD38* and *CD157* gene expression. Initial t-tests show that females are in the mean approximately 10 months younger than males (p < 0.001), likely owing to the fact that males are required to serve in the military before entering university. There are significant gender differences for IPS (p < 0.02), RPM (p = 0.004) and *CD157* expression (p=0.008). Males score higher on IPS than females, as previously reported (67). Similarly, a meta-analysis of RPM shows that among adults, the male advantage is 0.33 SD units (68). These sex differences in RPM appear likely to be influenced by sex differences in spatial ability (69). There are no significant differences between sex and measures of AUT, Openness, and *CD38* expression.

**Table 1.** Descriptive statistics for participants in the current study.

| Sex |  | CT | AUT | IPS | Openness | RPM | CD38 | CD157 | Age |
| --- | --- | --- | --- | --- | --- | --- | --- | --- | --- |
| Male | *M* | 16.74 | 15.19 | 1.55 | 162.61 | 7.16 | 11.00 | 6.66 | 22.20 |
|  | *SD* | 11.92 | 11.65 | 1.45 | 20.29 | 1.66 | 1.76 | 1.51 | 1.20 |
| Female | *M* | 15.55 | 14.68 | 0.87 | 163.51 | 6.44 | 10.42 | 5.75 | 21.32 |
|  | *SD* | 10.52 | 10.35 | 1.01 | 16.59 | 1.78 | 2.36 | 2.21 | 1.03 |
| Total | *M* | 16.16 | 14.94 | 1.22 | 163.05 | 6.81 | 10.71 | 6.21 | 21.77 |
|  | *SD* | 11.23 | 11.00 | 1.29 | 18.51 | 1.75 | 2.09 | 1.93 | 1.20 |

**Legend to Table 1.** Mean (*M*) and standard deviation (*SD*) of Creative Thinking, NEO-Openness, Raven's Progressive Matrices (RPM), age, and logged normalized values of *CD38* and *CD157* gene expressions. Creative Thinking (CT) scores are summations of Alternate Use Task (AUT) and Insight Problem Solving (IPS) tasks. $N=145$ .

### Association between creativity scores, RPM and Openness

We first examine associations between CT, fluid intelligence and personality (Figure 1). As expected, there is a correlation between CT and RPM, as well as with the personality trait of Openness, and the correlations are in the expected direction. Openness is correlated with total scores on CT in the pooled male and female sample. However, when the correlations are inspected in each of the two subscales, AUT and IPS, sex as we predicted tempers the strength of these correlations. Notably, AUT and IPS are not correlated (r=0.12, p > 0.1), confirming the orthogonal relationship between AUT and IPS in the multifaced landscape of creativity.

**Figure 1.**
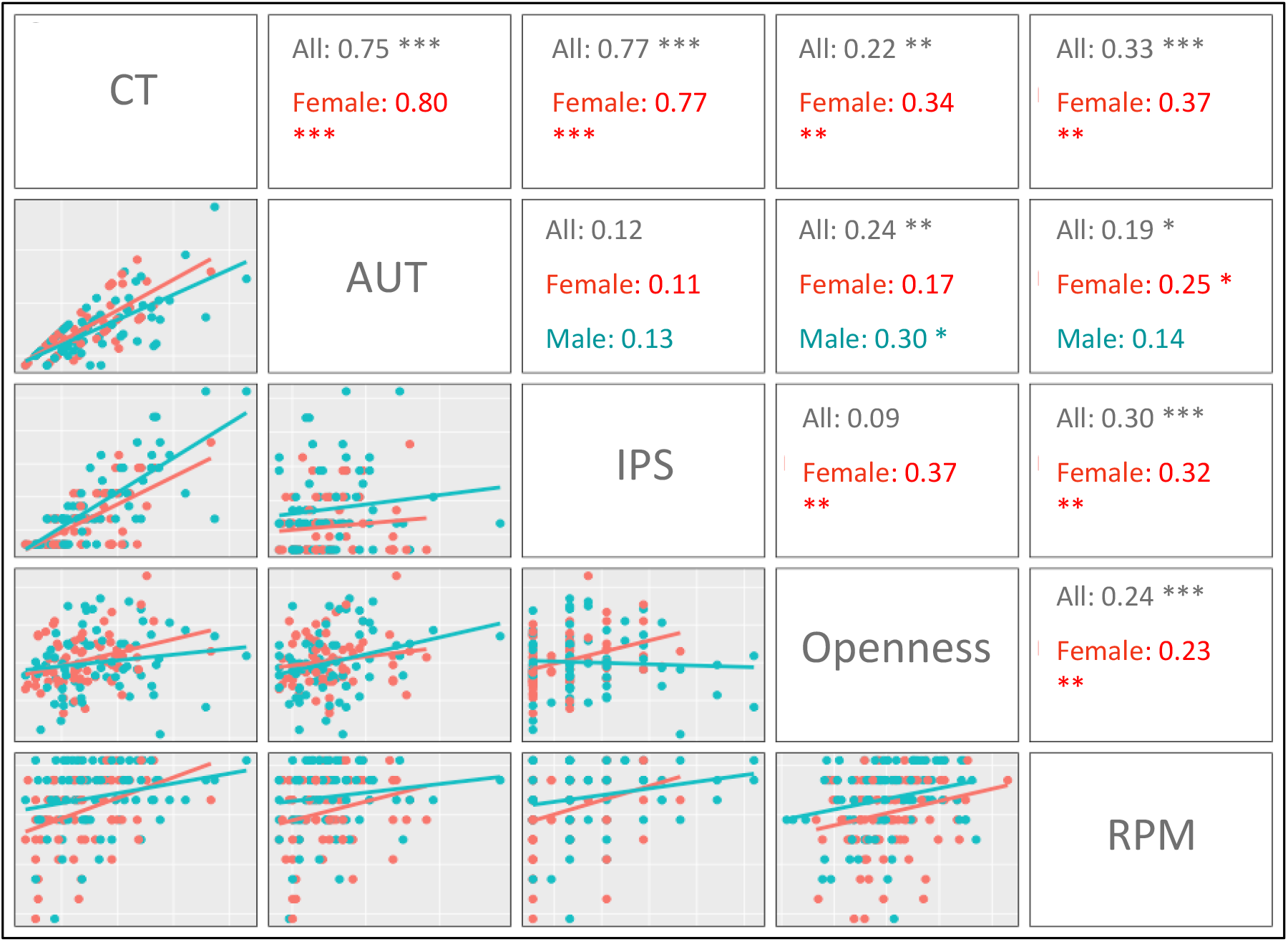
Scatterplots (lower triangle) and correlations (upper triangle) between CT, its components, AUT and IPS, with RPM and Openness. Significance: p < 0.001 ***, p < 0.01 **, p < 0.5 *.

### Relationship Between Creative Thinking (CT) and ADP-Ribosyl Cyclase Gene Expression

To test the association between *CD38*, *CD157* gene expressions and their interaction (focal independent variables) and total CT scores (the dependent variable), we carry out linear regression analyses (OLS) with robust standard error including age and sex in the model (Table 2, Model 1) which reached statistical significance (F (8,136) = 4.03, p < 0.001). Specifically, age is significantly correlated with CT (b=0.21, p = <0.05) while there is no main effect of *CD38* expression, however, a marginally significant main effect of *CD157* (b = −5.81, p = 0.08) is observed.

**Table 2.** The Relationship between Oxytocinergic Gene expression, Openness and Fluid Intelligence on Creative Thinking

| Variables | CT |  |  |  | AUT |  |  | IPS |  |  |
| --- | --- | --- | --- | --- | --- | --- | --- | --- | --- | --- |
|  | Model 1<br>All | Model 2<br>Male | Model 3<br>Female | Model 4<br>Female | Model 5<br>All | Model 6<br>Male | Model 7<br>Female | Model 8<br>All | Model 9<br>Male | Model 10<br>Female |
| CD38 | -3.51<br>(6.49) | -3.49<br>(6.50) | 2.90**<br>(1.31) | 3.56***<br>(1.15) | -8.35*<br>(4.93) | -8.41*<br>(4.91) | 1.73<br>(1.71) | 6.27<br>(3.95) | 6.17<br>(3.95) | 1.83*<br>(1.02) |
| CD157 | -5.81*<br>(3.30) | -5.75*<br>(3.35) | 8.89***<br>(3.07) | 8.51***<br>(2.65) | -5.37**<br>(2.64) | -5.32**<br>(2.63) | 5.02**<br>(1.99) | 0.65<br>(2.84) | 0.07<br>(2.84) | 3.48*<br>(1.76) |
| CD38 x CD157 | 8.18<br>(5.28) | 8.13<br>(5.30) | -9.60***<br>(3.32) | -9.45***<br>(2.87) | 9.13**<br>(4.46) | 9.03**<br>(4.45) | -5.64**<br>(2.25) | -2.84<br>(3.95) | -2.07<br>(3.94) | -3.81*<br>(1.99) |
| CD38 x sex | 6.37<br>(6.62) |  |  |  | 10.20*<br>(5.24) |  |  | -4.61<br>(4.14) |  |  |
| CD157 x sex | 14.67***<br>(4.52) |  |  |  | 10.43***<br>(3.39) |  |  | 2.94<br>(3.54) |  |  |
| CD38 x CD157 x sex | -17.74***<br>(6.24) |  |  |  | -14.76***<br>(5.07) |  |  | -1.17<br>(4.62) |  |  |
| sex | -3.81<br>(4.38) |  |  |  | -6.36*<br>(3.58) |  |  | 3.07<br>(2.82) |  |  |
| Age | 0.22**<br>(0.11) | 0.20<br>(0.14) | 0.24<br>(0.16) | 0.26**<br>(0.13) | 0.16**<br>(0.07) | 0.21**<br>(0.08) | 0.07<br>(0.11) | 0.10<br>(0.07) | 0.04<br>(0.10) | 0.19**<br>(0.08) |
| Openness |  |  |  | 0.02**<br>(0.01) | 0.01**<br>(0.00) | 0.01**<br>(0.00) | 0.00<br>(0.01) | 0.00<br>(0.00) | -0.01<br>(0.01) | 0.01***<br>(0.00) |
| RPM |  |  |  | 0.23***<br>(0.07) | 0.10**<br>(0.04) | 0.08<br>(0.05) | 0.13*<br>(0.06) | 0.15***<br>(0.04) | 0.18**<br>(0.07) | 0.10**<br>(0.05) |
| Constant | -3.12<br>(4.75) | -2.76<br>(5.17) | -7.51**<br>(3.57) | -12.96***<br>(3.27) | -0.33<br>(3.47) | -1.75<br>(3.49) | -4.20<br>(2.92) | -7.82**<br>(3.11) | -5.40<br>(3.67) | -8.76***<br>(2.04) |
| Observations | 145 | 74 | 71 | 71 | 145 | 74 | 71 | 145 | 74 | 71 |
| R-squared | 0.131 | 0.075 | 0.138 | 0.330 | 0.181 | 0.227 | 0.143 | 0.178 | 0.103 | 0.283 |

**Legend to Table 2.** Control variables used are age, sex, Openness and RPM. \*\*\* $p < 0.01$ , \*\* $p < 0.05$ , \* $p < 0.1$

### The Role of Sex in Moderating the Relationship Between CT and ADP-Ribosyl Cylase Gene Expression

Confirming our sex-differential hypothesis the analysis reveals a highly significant 2-way *CD157* x sex interaction (b = 14.67, p = 0.001) and a highly significant 3-way *CD38* x *CD157* x sex interaction effects(b = −17.74, p = 0.005) (details shown in Table 2, Model 1). Hence, we subsequently conduct separate regression analyses for males and females. Stratifying the participants according to sex (Table 2, males, Model 2; females, Model 3) reveals a significant association of total CT scores with ADP ribosyl-cyclase expression in females only. In females *CD38* (b = 2.90, p = 0.03), *CD157* (b = 8.89, p = 0.005), and their interaction *CD38* x *CD157* (b=-9.60, p = 0.005) reached significance. Notably, *CD38* and *CD157* expression are positively correlated with CT for females consistent with findings that the OT signalling in central pathways can enhance CT (15).

### Relationship Between CT, ADP-ribosyl cyclase expression, RPM and Openness to Experience

As others have observed there is a significant association between RPM, Openness and CT and for this reason we examine both these variables in our own data set. We test whether the correlation between CT and gene expressions is robust to inclusion of RPM and Openness. We conduct linear regression analysis of the full model (Table 2, Model 4) in females. The results remain robust to inclusion of Openness and RPM: *CD38* (b = 3.56, p = 0.003), *CD157* (b = 8.51, p = 0.002) and their interaction (b = - 9.45, p = 0.002). For females, the control variables Openness to experience (b = 0.02, p < 0.02), RPM (b = 0.23, p = 0.003) and age (b = 0.26, p < 0.05) are individually correlated with CT and account for approximately 20% of the variance in CT. Comparing Models 3 and 4 (Table 2), we observe that *CD38* and *CD157* gene expression as well as their interaction, are estimated to account for approximately 14% of the variance independent of Openness and RPM. Altogether, the total variance in female scores on total CT scores in the full model (Model 4), which includes oxytocinergic gene expression, Openness and RPM, explains a striking 33% of the variance.

### Relationship Between Creative Thinking Subscale (AUT and IPS) Analyses and Oxytocinergic Gene Expressions

Linear regression analyses of the subscales, AUT and IPS (dependent variables), for *CD38*, *CD157* gene expression, their interaction (independent focal variables) and the set of controlling variables (age, RPM and Openness to experience), are shown in Table 2 (Models 5-10). The detailed analysis is presented in the Supplementary Online Material. In brief, for AUT there is a main effect of *CD157* and a marginal effect of *CD38* and a significant *CD38* x *CD157* interaction. Again as predicted, sex tempers the relationships. For IPS, there is no main effect of ADP-ribosyl cyclases albeit it there are significant sex effects.

### Relationship Between Creative Thinking (CT, AUT and IPS) and Dopaminergic Gene Expressions

In the first stage of our analyses presented above, the focus is on the oxytocinergic pathway. Table 2 demonstrates that *CD38* and *CD157* significantly correlate with CT especially in females. Since considerable evidence links the mechanism of oxytocin action with direct action on dopaminergic pathways (70–72) and additionally, there is a known role of DA in creative cognition (24), we next examine *DRD2* and *COMT* gene expression on CT. First, we examine the relationship between *COMT* and *DRD2* on CT. As shown in Table 3, there is a significant correlation between both *DRD2* (b=2.9, p= 0.02) and *COMT* (b= −2.5, p=0.023) gene expression on CT scores in females (Table3, Model 3). *DRD2* is also significantly correlated to AUT (Table 3, Model 6: b=1.66, p=0.03) and IPS (Table 3, Model 9: b=1.24, p=0.04) for females. Notably, the coefficients are positive for *DRD2* suggesting that greater *DRD2* receptor expression, indicating higher DA tone, is as expected associated with higher CT scores. In addition, the coefficient for *COMT* is negative suggesting that metabolism of DA is as expected correlated with lower CT scores consistent with the results with *DRD2* receptor transcription. Overall these results point to an important role of dopaminergic activity on augmenting CT scores, mediated by a dual mechanism of increasing DA receptor number as well as reducing DA metabolism.

**Table 3.**
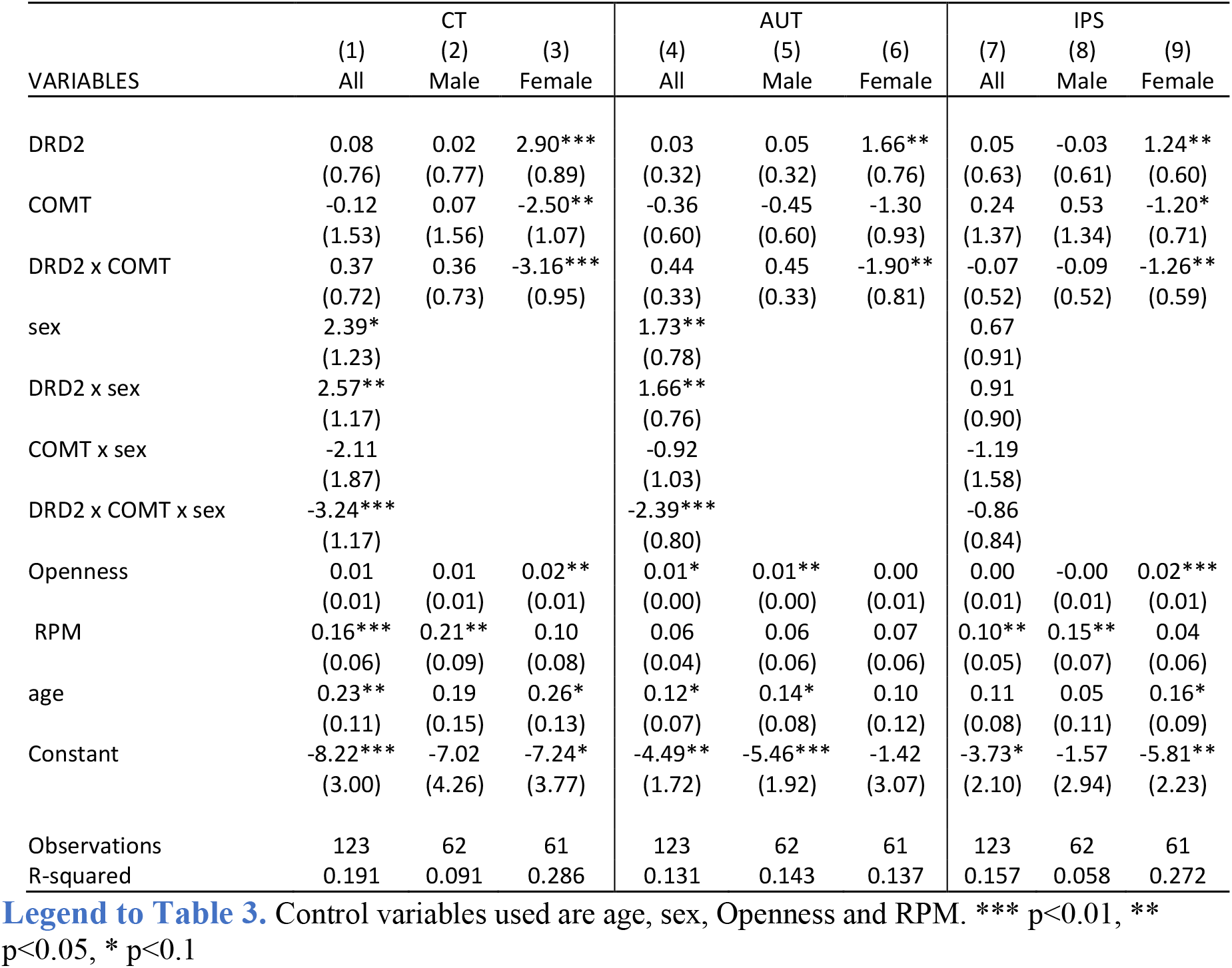
The Relationship between Dopaminergic Gene Expression, Openness, Fluid Intelligence and Creative Thinking

### Relationship Between CT and subscales, Oxytocinergic and Dopaminergic Gene Expressions, RPM and Openness to Experience

Finally, in the full model we include both oxytocinergic and dopaminergic expression, Openness and RPM. The model explains ∼39% of the variance in females (Table 4, Model 4). Gene expression by itself excluding Openness and RPM, explains ∼28% of the variance of CT (Table 4, Model 3). In Table 4 Models (5-10) show the results for the subscales of CT, AUT and IPS. In females the full model (gene expression, Openness, RPM and age) explains 22% of the variance for AUT and 35% of the variance for IPS. There are no correlations in male subjects with the sole exception of RPM.

**Table 4.**
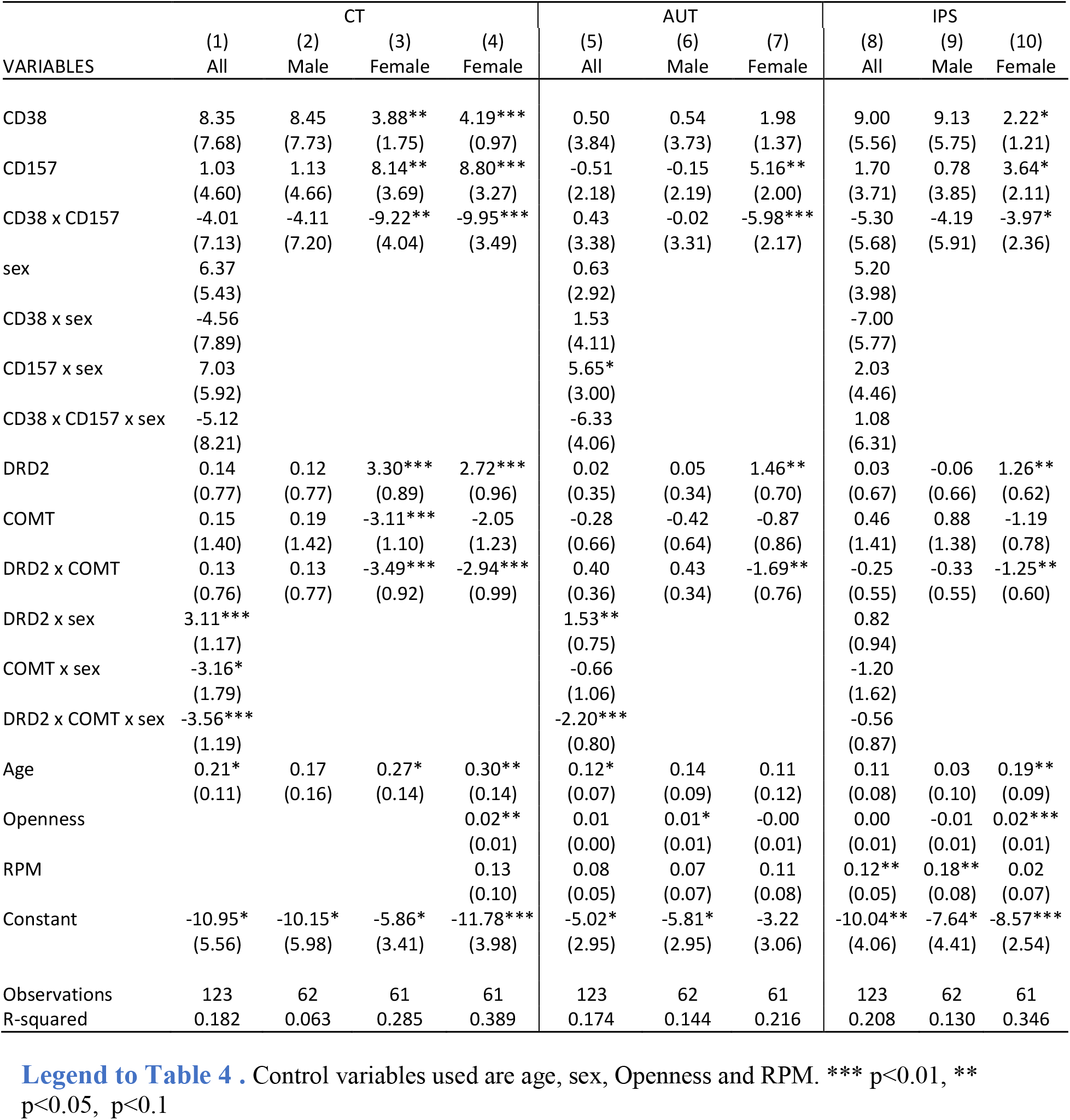
The Relationship between Oxytocinergic and Dopaminergic Gene expression, Openness and Fluid Intelligence on Creative Thinking

## Discussion

Recent findings from neuroimaging studies examining the neural signatures of CT using brain imaging including fMRI, PET and EEG (each method with its own strengths and weaknesses, e.g. (73)) have contributed to mapping the underlying brain pathways. Accumulating evidence points to the involvement of specific circuits and neural systems in the different components of the creative thinking process especially the medial prefrontal cortex (mPFC), the dorsolateral prefrontal cortex (DLPFC) and the dorsal anterior cingulate cortex (dACC) (74). The DLPFC and dACC are known to be activated across various domains of CT such as piano improvisation (39) creative story generation (75), word association (76), divergent thinking (76), fluid analogy formation (77), insight problem solving (77), and visual art design (77). Notably, OT, the paramount human social hormone (78), is a salient neuromodulator in these same brain regions prominent in CT. The actions of OT as a neuromodulator in the same brain regions identified with CT, focused our attention on this nonapeptide as a likely candidate underpinning human creativity.

Accumulating evidence also points to an important role for dopamine neurotransmission in CT. Dopamine pathways are widely distributed in the brain, including striatal pathways spanning the entire frontal cortex, and this neurotransmitter is shown in a plethora of investigations to be implicated in motivation, emotion, personality traits, and cognitive functions that are all related directly and indirectly to creativity (79). Genetic association studies, albeit characterized by small samples, more recently strengthen the role of dopaminergic neurotransmission in CT (22, 24). A rich literature attests to the close relationship between DA and OT brain pathways and their joint role in often underpinning the same complex behaviors (70–72, 80). Notably, oxytocin neuronal fibres impinge on DA cell bodies in the ventral tegmental area and oxytocin neurons also innervate the PFC, a target of dopaminergic input (81). Hence, it makes sense in the current investigation to examine along with the oxytocinergic system, the dopaminergic underpinnings of CT towards a more complete understanding of this salient human trait.

A further fine-grained approach to decipher the neurobiological underpinnings of CT, is to identify specific neurotransmitter pathways contributing to this trait. In humans a useful strategy towards this end, is to dip into the toolbox of molecular genetics. Despite the considerable heritability attributed to explaining individual differences in CT (3), only a few candidate gene association studies have been undertaken to identify genetic loci (15, 82). Moreover, gene association studies that only examine structural variants in the DNA code can account for only a very small portion of the variance of complex traits since environmental influences are ignored and the effect size of single SNP variations revealed by GWAS studies is quite small, e.g. educational attainment (65). A complementary strategy gaining traction that can identify jointly genetic and epigenetic contributions to complex behaviors such as CT is to implement an OMICS strategy using peripheral biomarkers, e.g. (83). We adopt this approach and examine gene expression in saliva along with measurements of salient psychological variables.

We find that the composite CT score (summing AUT and IPS) is significantly and positively associated with oxytocinergic (*CD38, CD157*) and dopaminergic (*DRD2*, *COMT*) gene expression only for females. When each of the two tasks that comprise the composite score is separately examined, we find *CD157* and *DRD2* expression is significantly correlated with AUT in females. There is only a weak correlation for IPS for the combined group as well as for each sex analysed separately. Sex effects of OT on behavior are frequently observed. For example, using a forced-choice affective body expression classification task, higher endogenous OT plasma levels were associated with better total recognition in both schizophrenic patients and controls and this association was specific to female patients (84). In a PET study using [^11^ C]raclopride that binds to dopamine receptors, subjects are exposed to a physical and emotional stressor (72). Female *OXTR* rs4813625 “C” allele carriers demonstrated greater stress-induced dopamine release, measured as reductions in receptor availability from baseline to the pain-stress condition relative to female “GG” homozygotes. Among the factors influencing individual differences in response to nasally administered OT, sex plays a salient role along with hormonal status and genetic factors. A brain imaging study (85) found that in subjects engaging in a DT task, their declarative memory related regions were strongly activated in men, whereas regions involved in theory of mind and self-referential processing were more engaged in women. Brain areas related to semantic cognition, rule learning and decision making were preferentially engaged in men during an AUT, whereas women displayed higher activity in regions related to speech processing and social perception. Similar sex-sensitive results are also observed in animal studies (86). To better understand sex effects of OT, it should be considered that gonadal sex hormones (estrogen, progesterone and testosterone) and OT and vasopressin (a closely related nonapeptide) co-evolved during the course of the development of mammalian affiliative behaviors and hence context-dependent actions of these nonapeptides on one or the other sex is not unexpected (87). Overall, there is extensive evidence indicating that at least some of oxytocin’s actions are sex specific, in some measure likely mediated by gonadal hormones.

Insight tasks evaluate an individual’s ability to resolve questions that require so-called cognitive restructuring of the problem representation (88). Insightful solutions look to be rapid cognitive events, the classical “Aha!” flash, albeit such ‘Aha!’ events may be prolonged and taking place in the unconscious (89). Notably, insightful problem solving often results in a single or at most very limited number of solutions. Indeed, in the current study only a single solution on the IPS is possible. In contrast, DT tasks markedly differ from ISP and are characterized as often only vaguely defined, viz. alternative uses for a button, and have open-ended solutions. The goal is for the subject being tested to produce several ideas and to seek especially original alternative solutions. DT tasks are generally scored for originality that captures the number of unique or rare ideas as well as fluency. Importantly, DT tasks are characterized by their reliability and have reasonable predictive validity (90). An investigation by Runco et al finds that DT is significantly correlated to certain creative activities after half a century (91). Fluency is a somewhat more difficult concept to assess than originality or uniqueness as it involves putting ideas into conceptual categories. A high correlation of (p=.89) between the latent fluency and originality variables is observed (92) and hence it is unlikely they convey distinct information (93). Overall, originality or uniqueness is considered as more central for creativity than fluency (33). Finally, originality has quite reliable variance, notwithstanding when the overlap with fluency is controlled for (94). For these reasons, in the current study we examined solely the originality facet of DT that appears to capture the core constituent of DT.

We took a novel approach to explore the neurochemical gene pathways underpinning creativity by implementing a partial OMICS strategy. Peripheral transcriptome studies (95) have gained momentum in many investigations. Interestingly, steady-state gene peripheral transcription appears to be partially heritable with an accounted variance *h*^*2*^ of ∼35% (96, 97). The modest heritability of peripheral transcription underscores that gene expression potentially captures not only structural variations in the genetic code but also a significant portion of environmental / epigenetic influences (the variance remaining when *h*^*2*^ is subtracted). The current investigation is one of the first to examine the neurobiological and neurochemical gene pathways in normal human cognition using a peripheral transcriptome approach and provides proof of principle that this approach provides a valuable perspective in understanding complex behaviors. We suggest the notion that it opportune for a paradigm shift in neurogenetic studies of complex behavioral traits towards a more inclusive OMICS strategy that also includes study of peripheral gene expression.

## Materials and Methods

### Participants

200 students (101 females, *M*_age_=21.82, *SD*=1.2) of Han Chinese descent from National University of Singapore were recruited via online advertisement in campus-wide student forum and were reimbursed $20 for their participation. Each one participant gave informed consent approved by the Institutional Review Board of National University of Singapore. Only participants with complete set of data for the Creativity measures of Alternative Uses Task (AUT) and Insight Problem Solving (IPS) task, NEO-PI-3 Openness Scale, Ravens Progressive Matrices (RPM, 9-item), CD38 and CD157 expressions were included in this study. The final number of participants included in this study is 145 (71 females).

### Questionnaires

#### Creativity Assessments

Creativity was assessed using the Alternative Uses Test (AUT) and the Insight Problem Solving (IPS) administered in a randomized order. AUT (98) measures divergent thinking and comprises of four measures of divergent thinking specifically originality, fluency, flexibility and elaboration. For the purpose of this study, we scored only for the originality component i.e. uses that are highly novel, that is an established indication of creativity (4, 99). Participants were instructed to give as many unconventional uses as possible for three objects randomly chosen for the list of objects used in AUT (button, nail and pencil) within 6 minutes for all three objects. Their responses were then scored against a separate norming sample of 72 similarly recruited NUS participants (not included in this study) to eliminate cultural bias that may be present for the objects. In the norming sample, the frequency of each use was calculated. Following Guilford’s scoring procedure (98) for AUT, a use that had frequency less than 2% were considered highly novel and given 2 points (e.g. using pencil to play pick-up-sticks). Uses with frequencies between 2-4% were given 1 point and uses that were given by more than 4% of the norming sample were given 0 point (e.g. using a nail to hang a picture on the wall). The final AUT score (M^AUT^ = 15.06, SD=11.2) was a summation of scores for all three objects.

Insight Problem Solving (IPS) task measures have often been linked to creativity and divergent thinking (98) as participants have to come up with novel solutions to uncommon problems that require suppression of self-imposed constraints acquired from previous experiences employing rules and boundaries in solving common problems. We tested participants ability to solve ten spatial insight problems from Dow and Mayer (29) (SI) within 10 minutes. Each correct answer was awarded 1 point while correct answers to problems that participants have encountered before were awarded half point to correct for task familiarity. The mean M_IPS_ = 1.26 and SD = 1.3.

For personality, participants were inventoried on their degree of openness to experiences using the Openness subscale of NEO-PI-3. The subscale examines facets such as fantasy, feelings, ideas, actions, aesthetics and values and consist of 48 items on a 5-point Likert scale. The mean score is 163.05 and *SD* = 18.50. Fluid intelligence was measured using the 9-item Abbreviated Form of the Raven’s Progressive Matrices (100) with mean 6.81 and *SD* 1.75.

### Saliva collection and Gene Expressions

#### Collection and Processing of Saliva

Samples were collected according to manufacturer’s protocol (DNA Genotek Inc., Ontario, Canada). Participants rinsed their mouth with clean water and abstained from food or drink for 1 hour before collection. About 2 mL of whole saliva deposited into the Oragene RE-100 tubes with preservative and stabilizer solution was derived by simulated habitual chewing for 2 mins. Sample was reacted with an in-cap stabilizer solution which was released upon cap replacement. The preserved saliva solution was neutralized in 1/25^th^ volume of Oragene RE-L2N solution on ice for 10 mins before clearance by centrifugation for 5 mins at 13000 RPM.

#### RNA extraction from Saliva

The extraction of RNA was performed using the RNeasy Micro kit (Qiagen, Hilden, Germany). Briefly, 500 ul of sample solution was mixed with 2 volumes of 95% ethanol and precipitated on ice for 30 min. The pellet was collected by centrifugation, completely dissolved in the RLT buffer to lyse the cells, then mixed with 70% ethanol. Nucleic acid was captured by centrifugation in spin column followed by in-column treatment with DNase I for 15 mins at room temperature. Sample was subjected through series of RPE buffer washes before final resuspension in 25 ul of RNase-free water. The quality and quantity of sample yield was assessed spectrophotometrically in the NanoDrop 2000 (Thermo Fisher Scientific, Waltham, MA).

#### Real-time RT-PCR analysis

Gene expression was determined by qPCR in a 2-gene custom RT2 Profiler PCR array system (Qiagen) according to manufacturer’s protocol. In the thermocycler, residual genomic DNA in 150 ng of total RNA sample was first treated in the genomic DNA elimination mix at 42°C for 5 mins, followed by the addition of reverse transcription mix and further incubation at 42°C for 15 mins. Template was added to the reaction mixture containing SYBR Green 1 Master mix with HotStart DNA polymerase which was then distributed to the PCR array at 25 ul per well. PCR was performed in the CFX96 qPCR detection system (Bio-Rad Laboratories, Hercules, CA) at cycling conditions that include an initial activation at 95°C for 10 min followed by 40 cycles of annealing and extension steps at 95°C for 15 seconds and at 60°C for 1 min respectively. A final default melting curve program was applied to generate a first derivative dissociation curve for each well. The expression levels were analysed using the ΔCT method from Cq values normalized to the expression of select reference from a series of candidate genes.

## Acknowledgements

Funding was provided by the AXA Research Fund, Templeton World Charity Foundation, and the Singapore Ministry of Education.

## Supplementary Material

### Subscale analysis of CT with oxytocinergic gene expressions, Table 2 Models 5-10

#### AUT

The associations between AUT and gene expression of *CD38*, *CD157*, and their interaction are significant when both males and females are jointly analysed (Model 5). There is a main effect of *CD157* (b = −5.32, p = 0.04) and a marginally significant effect of *CD38* (b = −8.35, p = 0.09). Their interaction is significant (b = 9.13, p = 0.04). In addition, there is a significant 2-way *CD157* x sex interaction (b = 10.43, p = 0.003) and *CD38* x sex interaction (b=10.20, p = 0.054) as well as a significant 3-way *CD38 x CD157* x sex interaction (b = - 14.76, p = 0.004). Separate analyses by sex show that gene expression (focal independent variable) is significantly correlated with AUT (dependent variable) for both males (Model 6) and females (Model 7). *CD157* expression is negatively correlated with AUT for males (b = −5.32, p = 0.047) and positively correlated for females (b = 5.02, p < 0.02). The *CD38* x *CD157* interaction is also significant for males (b = 9.03, p = 0.046) and females (b = −5.64, p = 0.015. *CD38* expression is marginally significant for males (b = −8.41, p <0.1) and not significant for females (p > 0.1). Openness to experience (b = 0.01, p = <0.05) and age (b = 0.22, p <0.05) are significantly correlated with AUT. For females, none of the control variables are significantly correlated with AUT.

#### IPS

For IPS (dependent variable), regression analysis conducted jointly with males and females is presented in Table 2 (Model 8). Only RPM is significant (b = 0.15, p < 0.001). and similar results are observed for males (Model 9), (b = 0.18, p = 0.013). For females (Model 10), IPS (dependent variable) is marginally correlated with *CD38* expression (b = 1.83, p = 0.077), *CD157* expression (b = 3.48, p = 0.052) and the interaction term is significant, *CD38 x CD157* (b = −3.81, p = 0.06). Each of the control variables is significantly correlated with IPS in females: Openness to experience (b = 0.01, p = 0.004), RPM (b = 0.1, p = 0.03) and age (b = 0.2, p = 0.02).

**Subscale analyses of CT, with dopaminergic gene expressions, Table 3, Models Subscale analyses of CT, with both oxytocinergic and dopaminergic gene expressions, Table 4, Models 5-10.**

